# Development and optimization of a novel T7 polymerase-independent Marburg virus minigenome system

**DOI:** 10.1101/2020.05.11.088211

**Authors:** Bert Vanmechelen, Joren Stroobants, Kurt Vermeire, Piet Maes

## Abstract

Marburg virus (MARV) is the only known pathogenic filovirus not belonging to the genus *Ebolavirus*. Minigenomes have proven a useful tool to study MARV, but all existing MARV minigenomes are dependent on the addition of an exogenous T7 RNA polymerase to drive minigenome expression. However, exogenous expression of a T7 polymerase is not always feasible and acts as a confounding factor in compound screening assays. We have developed an alternative minigenome that is controlled by the natively expressed RNA polymerase II. We demonstrate here the characteristics of this new system and its applicability in a wide range of cell types. Our system shows a clear concentration-dependent activity and outperforms the existing T7 polymerase-based system at higher concentrations, especially in difficult-to-transfect cell lines. In addition, we show that our system can be used for high-throughput compound screening in a 96-well format, thereby providing an attractive alternative to previously developed MARV minigenomes.

## Introduction

Marburg virus (MARV) is a highly pathogenic virus that was first discovered in 1967 in Germany (Siegert, Shu, Slenczka, Peters, & Muller, 1967; Smith, Simpson, Bowen, & Zlotnik, 1967). MARV is the only known member of the genus *Marburgvirus*, one of the five genera in the family *Filoviridae* (Amarasinghe et al., 2019). The best-known member of this family is Ebola virus (EBOV), which has caused two large outbreaks in the past few years and consequently has been the center of most filovirus-related research in the past decade (Rojas et al., 2020). Similarly to EBOV, MARV outbreaks are typically associated with a very high case-mortality rate. While advances have been made for Ebola virus disease in recent years, there are still no licensed therapeutics or vaccines available for Marburg virus disease (MVD)(WHO, 2018). Because of this lack of virus countermeasures, WHO has listed MVD as a ‘priority disease’ for research and development (Mehand, Al-Shorbaji, Millett, & Murgue, 2018). However, studies involving live MARV are difficult to perform due to its classification as a biosafety level 4 (BSL-4) pathogen (U.S. Department of Health and Human Services, 2009).

An alternative to handling infectious virus is the use of MARV minigenomes to study different aspects of the viral life cycle. In a minigenome system, the leader and trailer region of the genome are placed adjacent to a reporter gene and transfected into target cells alongside the different components of the viral polymerase complex (Wendt, Bostedt, Hoenen, & Groseth, 2019). Once the different viral proteins are expressed, they form polymerase complexes and bind to transcripts of the minigenome, leading to the expression of the reporter gene. The use of easily detectable reporters, such as luciferase or green fluorescent protein (GFP), in a BSL-2 setting, has enabled the widespread use of minigenome systems for viral research, although their applicability is usually limited to studying a specific phase of the viral life cycle, i.e., virus replication and/or transcription (Hoenen, Groseth, de Kok-Mercado, Kuhn, & Wahl-Jensen, 2011).

The first MARV minigenome was developed over 20 years ago, employing a chloramphenicol acetyltransferase as a reporter gene (Muhlberger, Lotfering, Klenk, & Becker, 1998). Since then, several variations of this minigenome have been made, aimed at improving its usability by replacing the reporter gene with different types of luciferases, GFP or even dual reporters, although the general design of the minigenome has been conserved (Schmidt & Muhlberger, 2016). In all currently existing MARV minigenome systems, the minigenome is placed immediately downstream of a T7 RNA polymerase promoter that controls the expression of the minigenome and ensures the generation of 5’ ends with minimal overhang. On the 3’ end, a self-cleaving hepatitis delta virus ribozyme (HdVRz) is attached to the minigenome to ensure the generation of defined 3’ ends, which are necessary for replication activity.

Although a potent T7 RNA polymerase promoter is now widely used to drive expression in viral minigenome systems, it does have certain drawbacks that limit its usability. As the T7 polymerase is derived from bacteriophages, it is not naturally expressed in mammalian cell lines and thus needs to be provided separately (Lieber, Kiessling, & Strauss, 1989). There are several ways to enable expression of a T7 polymerase in mammalian cells, such as co-transfection of an additional plasmid encoding for a T7 polymerase, or infection of cells with T7-expressing recombinant virus (e.g. replication-deficient vaccinia virus (MVA-T7))(Sutter, Ohlmann, & Erfle, 1995). However, plasmid-based expression of a T7 polymerase can be difficult to achieve in certain cell lines, while the use of MVA-T7 can lead to cytotoxicity, limiting the usability of T7 polymerase-driven systems in certain cell lines (Jordan, Horn, Oehmke, Leendertz, & Sandig, 2009; Nelson et al., 2017; Walpita & Flick, 2005). An example of such a cell line is the *Rousettus aegyptiacus*-derived RoNi/7.1 cell line. This is particularly troublesome when studying MARV, as *Rousettus aegyptiacus* bats are known to be the natural reservoir of MARV (Towner et al., 2009). A T7 polymerase can also act as a confounding factor in compound screening assays. Most minigenome systems used for antiviral compound screening are limited to detecting polymerase (co-factor) inhibitors. As such, the classes of compounds that are likely to show activity in these assays (e.g. nucleoside analogs) are also the ones that might inhibit T7 polymerase activity, potentially resulting in high rates of false positivity.

In this article, we describe the development and optimization of a MARV minigenome system under the control of a mammalian RNA polymerase II promoter. We show that this system is operational in a variety of cell lines and can be used as a high-throughput compound screening assay, providing a valid alternative for the widely used T7 polymerase-based systems.

## Results

### Development of a polymerase II-driven MARV minigenome

One of the key applications of minigenomes is the screening for viral polymerase inhibitors, highlighting the need for a system that is independent from exogenous polymerases. The necessary addition of such secondary polymerases, like the T7 polymerase, acts as a difficult-to-correct confounding factor. Conversely, in minigenomes using promoters targeting polymerases natively expressed in mammalian cell types, this confounding does not occur, as off-target inhibition of the host’s RNA polymerase will also directly affect cell viability. RNA polymerase II is the enzyme responsible for the generation of mRNA precursors in all eukaryotic cell types and as such provides an interesting alternative to the bacteriophage-derived T7 polymerase(Woychik & Hampsey, 2002). Furthermore, it has previously been shown that RNA polymerase II can be used to drive the expression of viral reverse genetics systems (Griffin et al., 2019; Martin, Staeheli, & Schneider, 2006; Nelson et al., 2017; Wang et al., 2015). To generate an RNA polymerase II-driven MARV minigenome, the trailer and leader regions of MARV, flanking an eGFP reporter gene, were cloned into the pCAGGS-3E5E-EBOV-eGFP vector, resulting in an antisense MARV minigenome under the control of the potent CAG promoter (Figure 1A). In addition, the trailer and leader regions of MARV were flanked by a self-cleaving hammerhead ribozyme and a HDV ribozyme, respectively, to ensure defined 5’ and 3’ ends. Transcription of this minigenome by the cellular RNA polymerase II and subsequent cleavage of the ribozyme sequences allows the minigenome to be bound by the viral polymerase (L) and its co-factors (NP, VP30 and VP35), which are provided in trans. The viral polymerase will then transcribe the reporter gene encoded by the minigenome, producing a measurable eGFP signal. Initial transfection tests showed that this CAG-controlled minigenome had only limited activity (∼25%) compared to the T7-driven MARV minigenome (Figure 1B). We hypothesized that the lack of minigenome activity (as evidenced by a weak GFP signal) was due to an incompatibility of the used hammerhead ribozyme with the MARV trailer region. As is the case for EBOV, the MARV trailer contains multiple conserved and self-complementary motifs, allowing the formation of secondary RNA structures. One of these structures is an RNA hairpin that is presumably recognized by the viral polymerase and acts as the replication promoter (Crary, Towner, Honig, Shoemaker, & Nichol, 2003; Volchkov et al., 1999). Because the formation of this hairpin is crucial for viral replication to occur, the incorporation of a flanking ribozyme sequence that generates defined 5’ ends containing the necessary motifs for secondary structure formation is indispensable to ensure minigenome activity and needs to be incorporated into the ribozyme design. As the genome ends of EBOV and MARV are highly conserved, we initially assumed we could reuse the hammerhead design used for EBOV in the MARV minigenome. This design is a type I hammerhead ribozyme, in which stem I is formed by pairing of the first eleven nucleotides of the virus trailer with a complementary motif at the 5’ end of the ribozyme sequence (Figure 2A). However, even though the first ∼20 nucleotides of the EBOV and MARV genomes are highly conserved, the upstream motifs differ, resulting in the formation of slightly different hairpin structures (Figure 2B). We hypothesized that these minor but distinct differences in structure and especially in the position of the EBOV and MARV trailer hairpins result in altered minigenome activity. Predictive modelling of RNA secondary structures in the EBOV trailer shows that a hairpin is formed at the very end (nt 1-54) of the sequence and that the addition of a hammerhead ribozyme, incorporating the first eleven nucleotides of the virus trailer, abolishes the formation of this hairpin. Conversely, the MARV hairpin is located slightly downstream (nt 12-62) and its formation is predicted to be unaffected by the addition of the ribozyme, thereby resulting in the attempted formation of two large secondary structures separated by only one nucleotide. Presumably, the presence of another large structure in the proximity of the hammerhead ribozyme results in sterical hindrance, preventing the ribozyme from being formed or cleaving itself, thereby yielding an inactive minigenome transcript.

**Figure 1.**
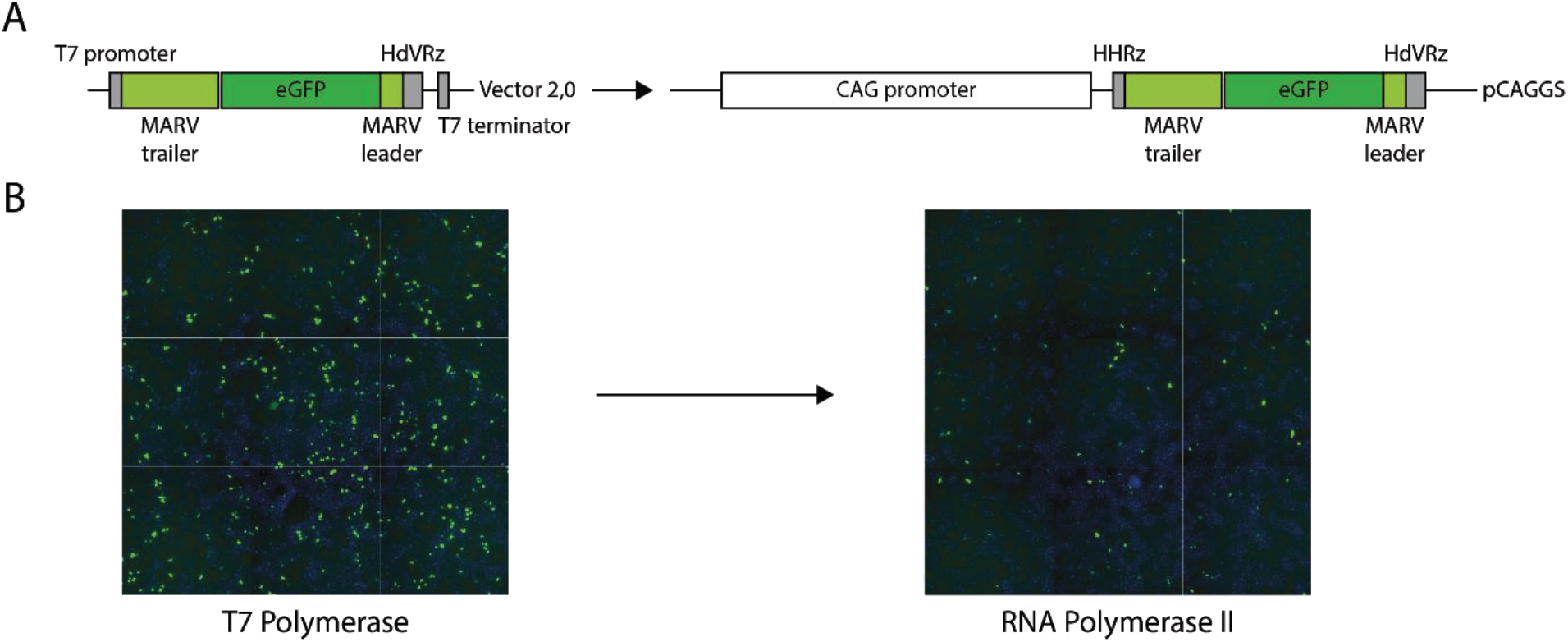
Exchange of the T7 promoter by a mammalian CAG promotor leads to a significant loss of MARV minigenome activity. (A) Schematic representation of the pCAGGS-HHRz-3M-eGFP-5M minigenome, made by replacing the T7 promoter of the T7-3M-eGFP-5M plasmid by a CAG promoter and a 5’ hammerhead ribozyme (HHRz). HdVRz = hepatitis delta virus ribozyme. (B) Comparison of the T7 polymerase and RNA polymerase II eGFP minigenome systems. HEK293T cells were transfected with 1000 ng of the indicated minigenome, as well as all necessary support plasmids. Images were taken 48 hours post transfection. Cells are stained with Hoechst 33342 as background staining (shown in blue), eGFP-expressing cells are shown in green.

**Figure 2.**
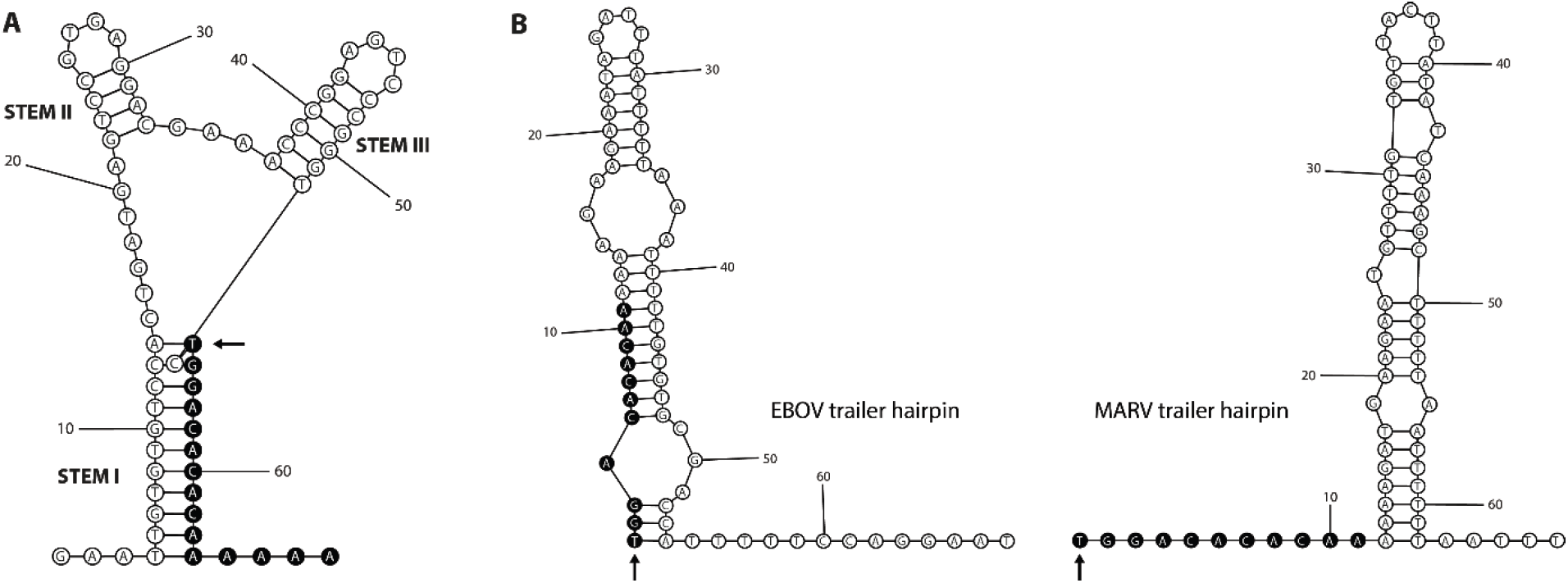
Secondary structure modelling of RNA structures. (A) Type I hammerhead ribozyme used in the initial pCAGGS-HHRz-3M-eGFP-5M minigenome. The arrow indicates the position at which the ribozyme cleaves. This corresponds to the start of the EBOV/MARV trailer, which is integrated into the 3’ end of the ribozyme sequence (marked in black) to ensure the generation of an exact 5’ end. (B) Comparison of the hairpin structures formed at the 5’ end of the EBOV and MARV trailers shows that the EBOV hairpin is positioned at the very end of the sequence while the MARV hairpin is shifted 11 nucleotides inwards. The start of each sequence is indicated with an arrow, while the residues marked in black correspond to the nucleotides incorporated into stem I of the hammerhead ribozyme.

To overcome the lack of minigenome activity, we sought to reduce this sterical hindrance by altering the relative size and position of the ribozyme and hairpin structures. However, simply adding a spacer sequence between the ribozyme and the MARV trailer hairpin to increase the distance between these two structures, or otherwise altering the sequence of the 3’ end of the ribozyme was not feasible, as the beginning of the MARV trailer forms part of stem I of the ribozyme. As such, modifications could only be made by mutating the 5’ end of the ribozyme sequence. Initially, we attempted to mimic the EBOV situation, in which the ribozyme and trailer hairpin share part of their sequence, resulting in the preferential formation of the hammerhead ribozyme. However, sequentially increasing the size of the first stem of the hammerhead to incorporate part of the hairpin sequence did not significantly increase minigenome activity (data not shown), presumably because the majority of the hairpin structure could still be formed. Because lengthening the stem of the ribozyme proved unsuccessful, we explored the opposite by shortening the stem as much as possible. The first eleven nucleotides at the 5’ end of the MARV trailer sequence, which represent one of the complementary strains that form stem I of the hammerhead ribozyme, do not belong to the trailer hairpin. Mutating the nucleotides at the 5’ end of the ribozyme that are complementary to these eleven nucleotides will therefore reduce base pairing in the ribozyme stem and consequently shorten the length of the stem without altering the MARV trailer sequence. As shown in Figure 3, each additional mutation that shortened the ribozyme stem and, consequently, increased the length of the spacer between the ribozyme and the trailer hairpin, resulted in an increase of minigenome activity (Fig 3A), with a maximum at a shortening of four nucleotides, corresponding to a remaining stem length of six base pairs. Further shortening the stem leads to a rapid loss of activity, in accordance with predictive RNA secondary structure modelling calculating the probability of the hammerhead structure being formed. Based on these findings, the six-basepair-stem hammerhead design was selected for further validation.

**Figure 3.**
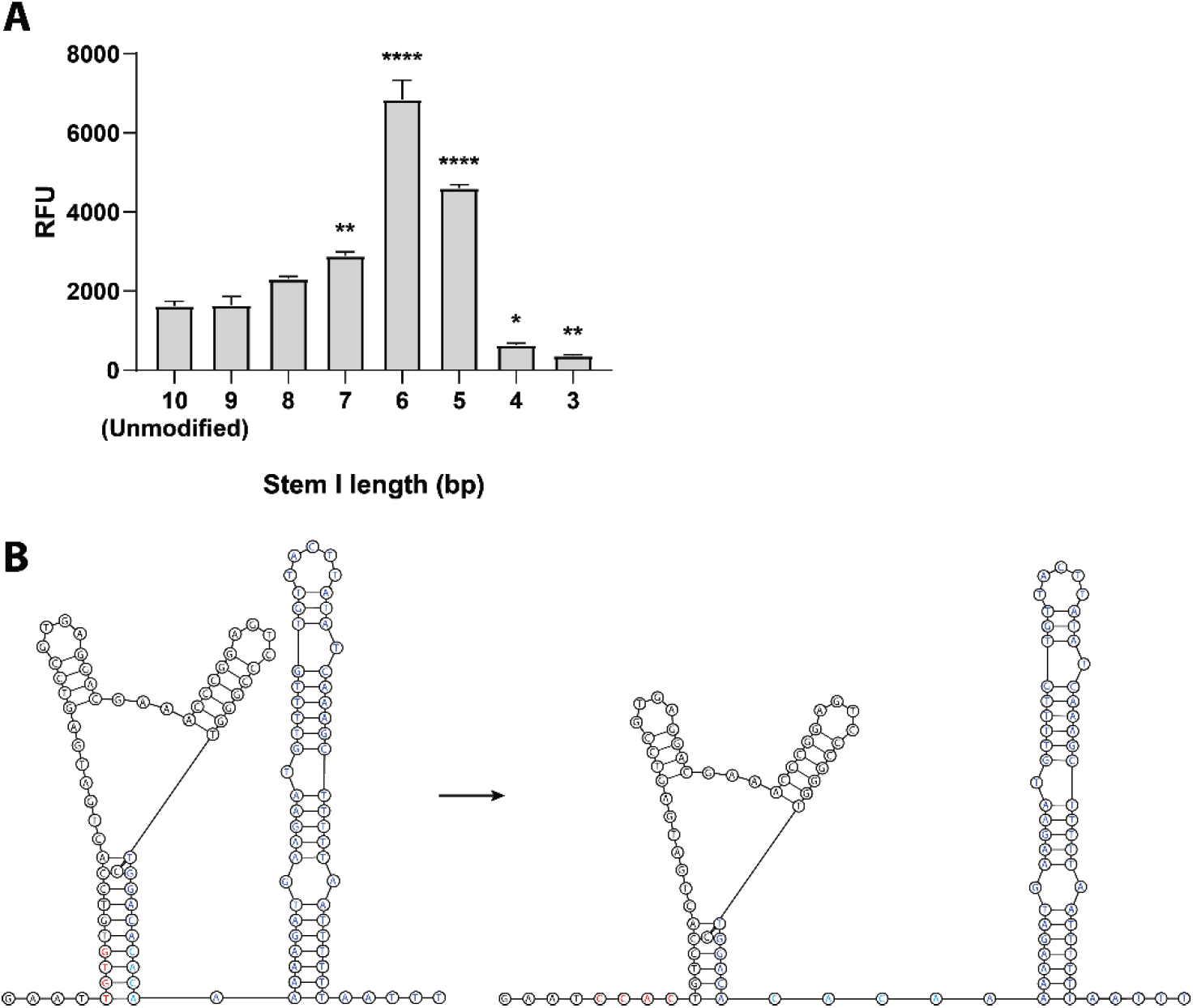
Shortening the ribozyme stem increases minigenome activity. (A) Transfection of HEK293T cells with different mutants of the pCAGGS-HHRz-3M-eGFP-5M minigenome shows that minigenome activity can be increased by reducing complementarity between the ribozyme and the MARV trailer. Shortening by 3-5 nucleotides significantly increases activity, with an optimum at a residual stem length of six base pairs. Data from three independent experiments, done in triplicate. Error bars indicate the standard error of mean. Significance testing was done using one-way ANOVA with Dunnett’s test for multiple comparisons: * p<0.05, ** p<0.01, **** p<0.0001. RFU = Relative fluorescence units, obtained by normalizing the amount of cells/well. (B) Schematic comparison of the unmodified (left) and the maximally active (right) ribozyme-trailer junctions. Nucleotides that are mutated to shorten the ribozyme stem are marked in red, their complementary partners that will act as additional spacers between the ribozyme and the MARV trailer in cyan and the rest of the MARV trailer in blue. See also supplementary table S1.

### Comparison of the T7 and RNA polymerase II MARV minigenomes

To qualitatively compare the optimized polymerase II-driven minigenome with the T7-polymerase control, HEK293T cells were transfected with MARV support plasmids and varying amounts of MARV minigenome plasmid. pCAGGS-T7 was added to all wells transfected with the T7 minigenome. Cells were incubated for 72 hours in an IncuCyte incubator, measuring cell density and fluorescence at distinct time points. As shown in Figure 4A, eGFP expression for both systems was almost undetectable after 24 hours but increased steadily over time. Both systems’ activity was dependent on the amount of minigenome transfected, with the T7 polymerase system reaching its optimum at lower concentrations (400 ng), while the polymerase II minigenome was more efficient when using higher amounts of plasmid (750/1000 ng). Using more than 1000 ng minigenome plasmid did not result in additional gain of activity (data not shown). Taken together, these data show that, at higher plasmid concentrations, the polymerase II-driven MARV minigenome outperforms the T7 polymerase system while displaying a similar time-dependent activity.

**Figure 4.**
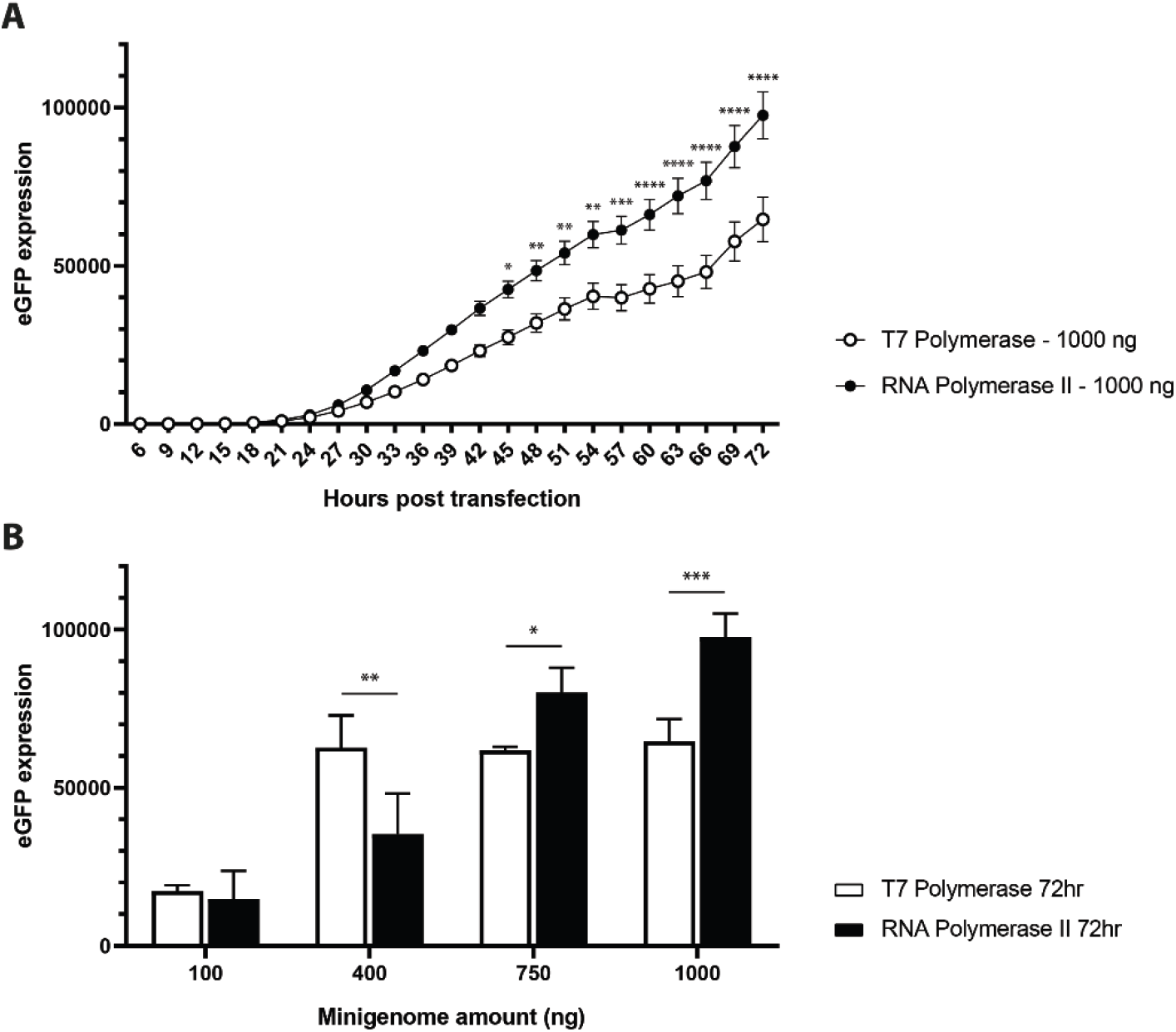
Comparison of the T7 polymerase- and RNA polymerase II-driven minigenome systems. (A) HEK293T cells were transfected with 1000 ng of T7 polymerase-or polymerase II-driven MARV minigenome and observed for 72 hours in an IncuCyte. Both systems displayed a similar time-dependent gain in eGFP expression, with the RNA polymerase II system reaching significantly higher expression levels. (B) While the T7 polymerase-driven MARV minigenome reaches its maximum activity at lower amounts of minigenome plasmid used, the RNA polymerase II-driven minigenome displays a clear concentration-dependent effect and outperforms the T7 system at higher plasmid concentrations. Data from three independent experiments, performed in triplicate in 6-well plates. Error bars indicate the standard error of the mean. eGFP expression is measured as the total green signal area, normalized for the amount of cells/well. Significance testing was done using multiple t-tests, correcting for multiple comparisons using the Holm-Sidak method: * p<0.05, ** p<0.01, *** p<0.001, **** p<0.0001. See also supplementary table S1.

To explore the cell dependency of the MARV minigenome system, we next compared the activity of the T7 polymerase and polymerase II-driven minigenomes in HEK293T cells and five additional cell lines (Figure 5). One of the drawbacks of T7 polymerase-dependent systems is that the T7 polymerase is not natively expressed in most mammalian cell lines and needs to be supplied either by co-transfection or through transduction with the MVA-T7 vaccinia virus. However, supplementing the transfection mix with an additional (T7) plasmid reduces transfection efficiency, while not all cell lines are equally suited for transduction. This is particularly true for *R. aegyptiacus* cell lines, which are known to die rapidly when transduced with MVA-T7 (Jordan et al., 2009). As *R. aegyptiacus* bats are the natural host for MARV, the availability of a minigenome that can be employed in these cell lines is a highly desired tool for MARV research. Therefore, in addition to cell types commonly used in varying types of filovirus research (Albarino et al., 2013; Enterlein et al., 2006; Tsuda et al., 2015), such as African green monkey Vero E6 cells, hamster-derived BHK21 and BSR-T7/5 cells and the human HEK293T and Huh-7 cell lines, we tested our system in the *R. aegyptiacus* RoNi/7.1 cell line, all. For this experiment, all cell lines were transfected with 1000 ng of either the T7 polymerase-or polymerase II-driven MARV minigenome, encoding a Renilla luciferase as reporter gene. After 72 hours, cells were lysed and luciferase activity measured. As summarized in Figure 5, the activity of the polymerase II system was comparable to or higher than the activity of the T7 system. Especially in the difficult-to-transfect Vero E6 and RoNi/7.1 cells, the polymerase II system clearly outperformed the T7 system. The only exception was the BSR-T7/5 cell line, where the T7 system yielded higher luciferase levels than the polymerase II system. This cell line is a modified clone of the BHK21 cell line that stably expresses T7 polymerase (Buchholz, Finke, & Conzelmann, 1999). Presumably, this native expression of T7 polymerase contributes to the increased activity of the T7 polymerase-driven minigenome in this cell line. Together, these results show that the polymerase II system can be successfully used in a wide variety of cell lines suitable for filovirus research, including cells from *R. aegyptiacus* bats, the natural hosts for MARV.

**Figure 5.**
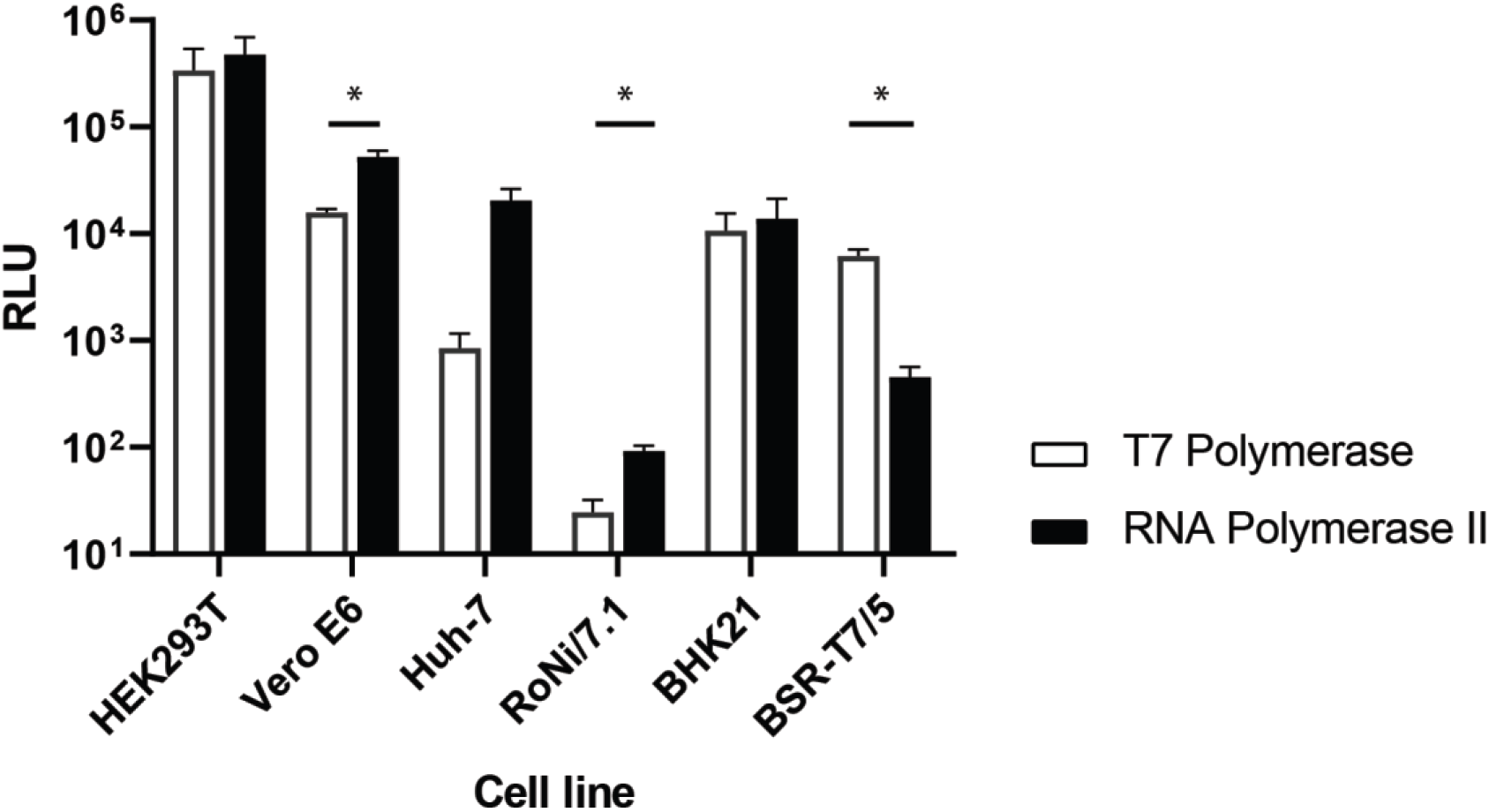
Minigenome activity comparison in different cell lines. Different cell lines were transfected with all support plasmids and 1000 ng of the T7 polymerase-driven or the modified RNA polymerase II-driven MARV minigenome containing a Renilla luciferase. Data is summarized from three independent experiments, performed in triplicate. RLU = Relative luciferase units, background-corrected. Significance testing was done using multiple t-tests, correcting for multiple comparisons using the Holm-Sidak method: * p<0.05. See also supplementary table S1.

### The polymerase II system as a tool for antiviral research

One of the primary applications of minigenome systems is their use in antiviral compound screening. Even though they can only detect polymerase (cofactor) inhibitors, their ease of use makes them an often sought-out alternative to working with live virus. To show that the polymerase II system presented here can be used for antiviral screening, HEK293T cells were transfected with the polymerase II minigenome and all pCAGGS MARV support plasmids. Six hours post transfection, cells were transferred to a 96-well plate prefilled with a dilution series of ribavirin. Ribavirin is a broad-spectrum antiviral compound that is used for the treatment of viral hemorrhagic fevers, respiratory syncytial virus infections and hepatitis C virus infections, although its use is not recommended for the treatment of flavivirus and filovirus infections due to a lack of proven clinical benefit (Huggins, 1989). Nonetheless, ribavirin possesses detectable *in vitro* anti-MARV activity in the polymerase II MARV minigenome system with an IC50 value of 23.7 µM (CC50: 90.9 µM), in accordance with previous results obtained using a T7-driven MARV minigenome (Figure 6)(Uebelhoer et al., 2014).

**Figure 6.**
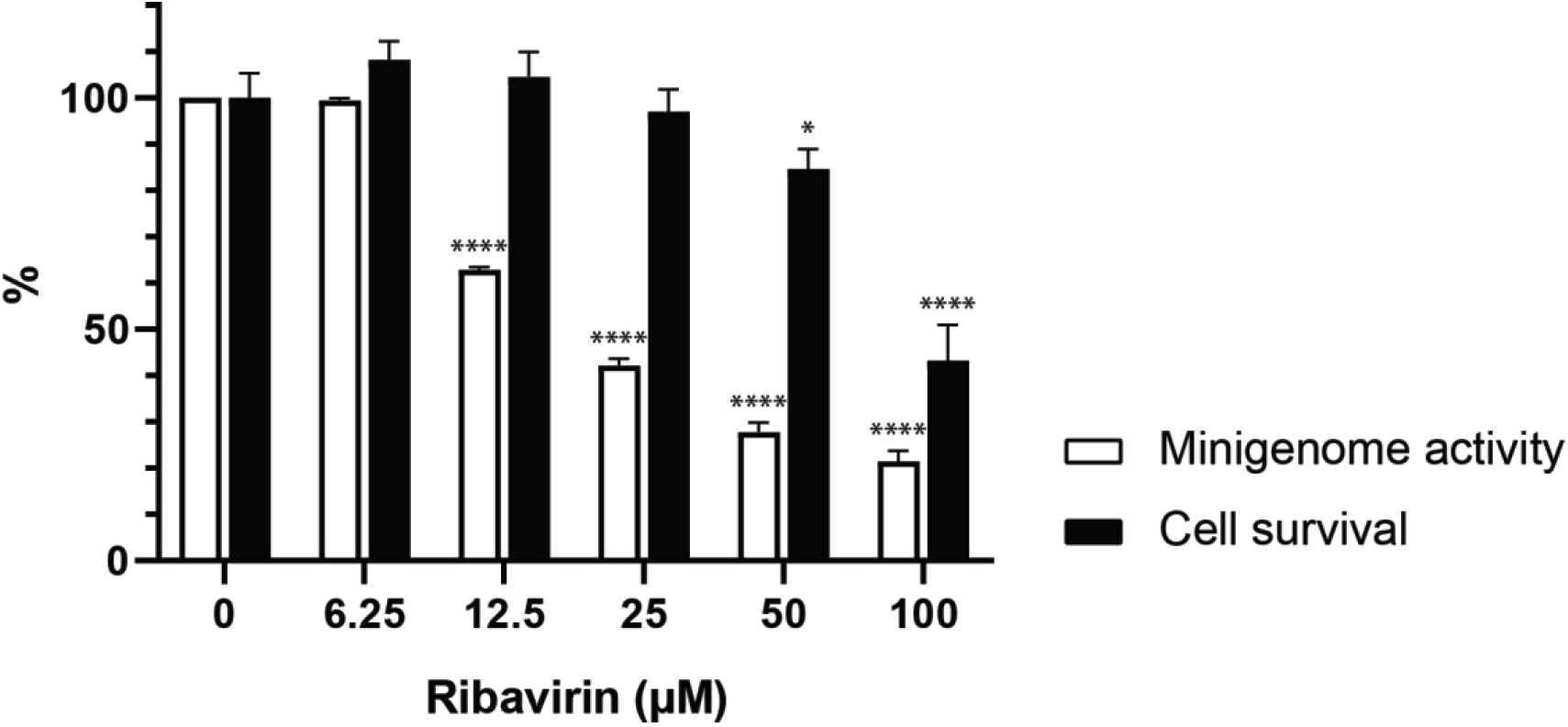
Ribavirin displays selective anti-MARV activity in vitro. HEK293T cells were transfected with the modified RNA polymerase II-driven MARV minigenome and, after 6 hours, transferred to 96-well plates prefilled with dilutions of ribavirin. After 48 hours, a clear dose-dependent reduction in minigenome activity was observed. Data from three independent experiments, performed in triplicate. Significance testing compared to the untreated control (0 µM) was done by two-way ANOVA with Dunnett’s multiple comparisons test: * p<0.05, **** p<0.0001. See also supplementary table S1.

To efficiently use a minigenome system for compound screening, scaling up is necessary to reduce the monetary and time cost per compound. By performing transfections in T-75 flasks, using the inexpensive PEI as a transfection reagent and subsequently transferring the transfected cells to 96-well plates filled with compound, well-to-well variation is minimized, increasing the reliability of the obtained results. Using this method, the polymerase II MARV minigenome system was shown to have a Z’ score of 0.8, indicating a highly robust assay (Zhang, Chung, & Oldenburg, 1999).

In conclusion, we present here a novel MARV minigenome system. By specifically tailoring the hammerhead ribozyme sequence to that of the MARV trailer, the minigenome could effectively be placed under the control of an RNA polymerase II promoter. The system shown here represents the first MARV minigenome that can work independently from the addition of an exogenous polymerase, allowing the system to be used in a high-throughput manner and in a wide range of cell lines while concurrently avoiding the lower specificity typically observed with T7 polymerase-based systems.

## Materials and methods

### Cell lines

Cell lines used for this study were human embryonic kidney cells (HEK-293T), African green monkey kidney cells (Vero E6; American Type Culture Collection, C1008), hamster kidney fibroblasts (BHK21, BSR-T7/5), human hepatocellular carcinoma cells (Huh-7) and kidney fibroblasts from the fruit bat *Rousettus aegyptiacus* (RoNi/7.1), kindly provided by Prof. M.A. Müller, Charité, Berlin, Germany. All cell lines were maintained in Dulbecco’s Modified Eagle Medium (DMEM; Thermo Fisher Scientific, MA, USA) supplemented with 10% fetal bovine serum (FBS; Biowest, France), with the exception of the BHK21 cells, which were maintained in Minimal Essential Medium (MEM) Rega-3 (Thermo Fisher Scientific) + 10% FBS. 200 mM L-glutamine (Thermo Fisher Scientific) was added to the BHK21, Huh-7, BSR-T7/5 and RoNi/7.1 cell medium. Additional supplements were 1% sodium bicarbonate (Thermo Fisher Scientific) for Huh-7 and BHK21 cells, 1% sodium pyruvate (Thermo Fisher Scientific) for RoNi/7.1 cells and 1% Non-Essential Amino Acids (NEAA; Thermo Fisher Scientific) for Huh-7 and RoNi/7.1 cells. Used antibiotics were 1% Penicillin-Streptomycin (Thermo Fisher Scientific) for Vero E6, BSR-T7/5, Huh-7 and RoNi/7.1 cells, 0.5% Geneticin (InvivoGen, France) for BSR-T7/5 cells and 1% Gentamicin (Thermo Fisher Scientific) and 0.2% Fungizone (Thermo Fisher Scientific) for Vero E6 cells.

### Minigenome design and construction

The artificial T7-3M-Luc-5M minigenome and plasmids encoding the MARV L, NP, VP30 and VP35 proteins were kindly provided by Prof. Stephan Becker. The T7-3M-eGFP-5M minigenome was made by replacing the Renilla luciferase gene in the T7-3M-Luc-5M minigenome with an eGFP gene through assembly of PCR fragments using the NEBuilder HiFi DNA assembly cloning kit (New England Biolabs, MA, USA). A modified version of this minigenome, containing a multiple cloning site and a hammerhead ribozyme (HHRz) sequence between the T7 promoter and MARV trailer sequence (T7-HHRz-3M-eGFP-5M), was made by mutagenesis PCR using the site-directed mutagenesis kit (New England Biolabs). Additional mutants of this intermediate vector were generated through site-directed mutagenesis of the HHRz sequence. Each of the resulting MARV minigenomes was then cloned into the pCAGGS_3E5E_eGFP vector, which was a gift from Elke Mühlberger (Addgene plasmid # 103054). SacI-HF (New England Biolabs) was used for restriction digestion, followed by vector-insert ligation using the Quick Ligation kit (New England Biolabs) to generate the original and mutated pCAGGS-HHRz-3M-eGFP-5M vectors. Sanger sequencing was used to confirm the sequence of all vectors. pCAGGS-3M-Luc-5M minigenomes were made using the same approach.

### Transfection and eGFP imaging

TransIT transfection reagent (Mirus Bio, WI, USA) and Lipofectamine LTX were used according to the manufacturer’s instructions. TransIT-293 was used for the transfection of HEK293T cells, TransIT-LT1 for the transfection of BHK-21J, Huh-7 and Vero E6 cells and Lipofectamine LTX for the transfection of BSR-T7/5 and RoNi/7.1 cells. One day prior to transfection, cells were seeded in 6-well plates at sufficient density to reach 80-90% confluence at the time of transfection. The next day, a mixture consisting of 3:1 (DNA:reagent) transfection reagent and 1000 ng pCAGGS-L, 500 ng pCAGGS-NP, 100 ng pCAGGS-VP35, 100 ng pCAGGS-VP30 and 1000 ng pCAGGS-HHRz-3M-eGFP-5M/pCAGGS-HHRz-3M-rLuc-5M or 1000 ng T7-3M-eGFP-5M/T7-3M-rLuc-5M and 1000 ng pCAGGS-T7 was added to each well. Six hours post transfection, cells were trypsinised and transferred to a 96-well plate (5 × 10^4^ cells/well), prefilled with 100 µl DMEM +2% FBS/well. When using rLuc as a reporter, plates were incubated at 37°C for 72 hours prior to assessing luciferase activity using the Renilla-Glo Luciferase Assay System (Promega, Madison, WI, USA). Readout of eGFP was done by incubating and monitoring plates at 37°C for 72h in an IncuCyte® (Essen BioScience Inc., Ann Arbor, MI, USA) for real-time imaging, or by incubating at 37°C for 48h and subsequently adding 6 µM Hoechst 33342 nucleic acid stain (Thermo Fisher Scientific) as a background stain for high content imaging analysis, performed on an Arrayscan XTI (Thermo Fisher Scientific). Fluorescence signal was generated by exciting the cells with the following wavelengths: 386-23 nm (Hoechst) and 485-20 nm (eGFP), and images were taken with a 10X objective. Exposure times and objective offsets were kept constant for each imaging plate. A custom analysis protocol was developed in-house using the Cellomics SpotDetector BioApplication. First, a background reduction was performed on both channels to ensure positive signals. Second, nuclei were detected using a dynamic tresholding method based on differences in intensity peaks. Next, eGFP signals were detected using a fixed-intensity threshold based on images acquired from the eGFP positive control. Additional parameters were determined to remove non-nuclei and non-eGFP objects (false positive signals). Lastly, the percentage and number of eGFP positive cells was calculated using the Cellomics software. For compound screening, transfections were done in T-75 flasks using polyethylenimine (Polysciences Inc, PA, USA). Transfections were done according to the manufacturer’s instructions, using 8 µg pCAGGS-L, 4 µg pCAGGS-NP, 800 ng pCAGGS-VP35, 800 ng pCAGGS-VP30 and 8 µg pCAGGS-HHRz-3M-eGFP-5M for each flask.

### RNA secondary structure modelling and statistical analyses

Secondary structure modelling of the hammerhead ribozyme and MARV/EBOV trailer hairpins was done using the RNAstructure Fold and MaxExpect web servers (https://rna.urmc.rochester.edu/RNAstructureWeb/). Comparisons of minigenome activity and associated statistical analyses were done using GraphPad Prism 8.2.0. Z-factor calculations for compound screening were done using the equation: Z = 1-[(3_σ_C_+_ + 3_σ_C_-_)/(_µ_C_+_ - _µ_C_-_)](Zhang et al., 1999). Positive control wells were transfected with all elements of the minigenome assay, while the pCAGGS-L plasmid was omitted from the negative controls.

## Acknowledgments

The authors wish to thank prof. E. Mühlberger, Boston University, MA, USA for providing the pCAGGS_3E5E_eGFP plasmid, prof. S. Becker, Philipps-Universität, Marburg, Germany for providing the T7-3M-Luc-5M minigenome and MARV support plasmids, and Prof. M.A. Müller, Charité, Berlin, Germany for providing the RoNi/7.1 cell line. The authors also wish to thank Winston Chiu and Leentje Persoons for excellent technical assistance. BV is supported by a FWO SB grant for strategic basic research of the “Fonds Wetenschappelijk Onderzoek”/Research foundation Flanders [1S28617N]. This work is also supported by ‘Interne Fondsen KU Leuven / Internal Funds KU Leuven’ project 3M170314 awarded to KV and PM. The funders had no role in study design, data collection and analysis, decision to publish or preparation of the manuscript.

## Declarations of interest

None

## Author contributions

BV, JS and PM conceived the experiments. BV and JS performed the experimental work. BV, KV and PM wrote the main manuscript. KV and PM supplied laboratory materials and funding. All authors revised and approved the manuscript.

## Supplementary data

**Supplementary table S1**

Detailed overview of the statistical test results obtained in figures 3, 4, 5 and 6.

## Source data

**Figure 3-source data**

Data used to create the graph in figure 3.

**Figure 4-source data A**

Data used to create the graph in figure 4A.

**Figure 4-source data B**

Data used to create the graph in figure 4B.

**Figure 5-source data**

Data used to create the graph in figure 5.

**Figure 6-source data**

Data used to create the graph in figure 6.

